# Carboxy-terminal blockade of sortilin binding enhances progranulin gene therapy, a potential treatment for frontotemporal dementia

**DOI:** 10.1101/2024.09.15.613118

**Authors:** Shreya N. Kashyap, Stephanie N. Fox, Katherine I. Wilson, Charles F. Murchison, Yohannes A. Ambaw, Tobias C. Walther, Robert V. Farese, Andrew E. Arrant, Erik D. Roberson

**Affiliations:** Center for Neurodegeneration and Experimental Therapeutics, Alzheimer’s Disease Center, Department of Neurology, University of Alabama at Birmingham; Birmingham, USA; Sloan Kettering Institute, Memorial Sloan Kettering Cancer Center; New York, USA; Howard Hughes Medical Institute, New York, USA

## Abstract

Frontotemporal dementia is commonly caused by loss-of-function mutations in the progranulin gene. Potential therapies for this disorder have entered clinical trials, including progranulin gene therapy and drugs that reduce progranulin interactions with sortilin. Both approaches ameliorate functional and pathological abnormalities in mouse models of progranulin insufficiency. Here we investigated whether modifying the progranulin carboxy terminus to block sortilin interactions would improve the efficacy of progranulin gene therapy. We compared the effects of treating progranulin-deficient mice with gene therapy vectors expressing progranulin with intact sortilin interactions, progranulin with the carboxy terminus blocked to reduce sortilin interactions, or GFP control. We found that expressing carboxy-terminally blocked progranulin generated higher levels of progranulin both at the injection site and in more distant regions. Carboxy-terminally blocked progranulin was also more effective at ameliorating microgliosis, microglial lipofuscinosis, and lipid abnormalities including ganglioside accumulation and loss of bis(monoacylglycero)phosphate lipids. Finally, only carboxy-terminally blocked progranulin reduced plasma neurofilament light chain, a biomarker of neurodegeneration, in progranulin-deficient mice. These results demonstrate that modifying the progranulin cargo to block sortilin interactions may be important for increasing the effectiveness of progranulin gene therapy.

**One-sentence Summary:** The effectiveness of progranulin gene therapy in models of FTD is improved by blocking the protein’s carboxy terminus, which prevents sortilin binding

## INTRODUCTION

Heterozygous loss-of-function mutations in the progranulin gene (*GRN*) account for 5– 20% of familial cases and up to 5% of sporadic cases of frontotemporal dementia (FTD), a relatively early-onset dementia characterized by social, behavioral, and language abnormalities (*1, 2*). Histopathological characteristics of FTD include tau or TDP-43 pathology (*3*), lipofuscinosis (*4, 5*), astrogliosis (*6*), microgliosis (*7*), and neurodegeneration (*8*). Homozygous loss-of-function mutations in progranulin cause a form of neuronal ceroid lipofuscinosis (CLN11), a lysosomal storage disorder histopathologically characterized by lipofuscinosis, neuroinflammation, and retinal degeneration (*9–12*). The overlapping pathology of FTD-*GRN* and CLN11 underscores the lysosomal function of progranulin which mediates both its neurotrophic properties (*13*) and its role in microglial activation (*14*). Of the more than 70 *GRN* mutations associated with FTD, nearly all cause progranulin haploinsufficiency (*1, 15–17*). Therefore, progranulin replacement, including progranulin gene therapy using adeno-associated virus (AAV) vectors, is a conceptually straightforward therapeutic approach for both FTD-*GRN* and CLN11 (*18*).

Progranulin gene therapy allows for specific and sustained restoration of brain progranulin, and multiple preclinical studies have demonstrated its therapeutic potential in progranulin-deficient mouse models of FTD-*GRN* and CLN11 (*19–24*). We previously found that intraparenchymal delivery of AAV1–mouse progranulin in the medial prefrontal cortex (mPFC) of *Grn^−/−^* mice corrects lysosomal enzyme abnormalities, including cathepsin D and β-glucocerebrosidase, and decreases lipofuscinosis and microglial pathology even in regions distant from the injection site, including hippocampus and thalamus (*20, 25*). We also found that intraparenchymal injection of AAV-*Grn* into the mPFC of *Grn*^+/–^ mice corrects social behavioral deficits (*19*). Others found similarly that intracerebroventricular delivery of AAVhu68–human progranulin boosts CSF progranulin levels and reduces lipofuscinosis and microgliosis in *Grn*^−/−^ mice (*22*). Building on this preclinical foundation, there are now three progranulin gene therapies in Phase 1/2 clinical trials (NCT04408625, NCT04747431, and NCT06064890).

Notably, in our previous studies showing a benefit of AAV-progranulin (*19, 20*), a tag was added to the progranulin C-terminus to allow identification of the exogenous gene therapy cargo. Importantly, the progranulin C-terminus is critical for its interaction with sortilin (*26, 27*) and the C-terminal blockade caused by this tag reduces sortilin binding (*20*). Nonetheless, progranulin efficiently localized to neuronal lysosomes *in vivo* and reduced lysosomal pathology both in neurons and microglia, indicating that the therapeutic effects of AAV-progranulin do not require sortilin interactions (*19, 20*). But because reducing sortilin interactions boosts progranulin levels and is the basis of the anti-sortilin antibody approach to treating FTD-*GRN* (*23, 26, 28, 29*), we asked if this C-terminal tag and resulting blockade of sortilin binding may actually enhance the efficacy of AAV-progranulin. Here, we address this question by directly comparing the effects of AAVs expressing C-terminally blocked progranulin versus progranulin with intact sortilin binding.

## RESULTS

### Blocking the progranulin C-terminus increases AAV-derived progranulin levels

To examine the effects of blocking the progranulin C-terminus on gene therapy effectiveness, we created an AAV expressing progranulin with a tag at the N-terminus after signal peptide cleavage (NT-*Grn*), leaving the C-terminus unblocked (Fig. 1A). We previously showed that this NT-*Grn* construct has intact sortilin binding, while the C-terminally blocked construct does not (*20*). Consistent with prior results with *Grn*-CT and the expected tropism of the vector, exogenous progranulin derived from both *Grn*-CT and NT-*Grn* was detected predominantly in neurons and not in microglia (fig. S1). We treated *Grn^−/−^* mice with AAV vectors carrying either GFP control, the previously used C-terminally tagged progranulin (*Grn*-CT), or NT-*Grn*. All vectors were packaged in an AAV1 capsid with AAV2 ITRs and delivered via bilateral intraparenchymal injections in the mPFC.

**Fig. 1.**
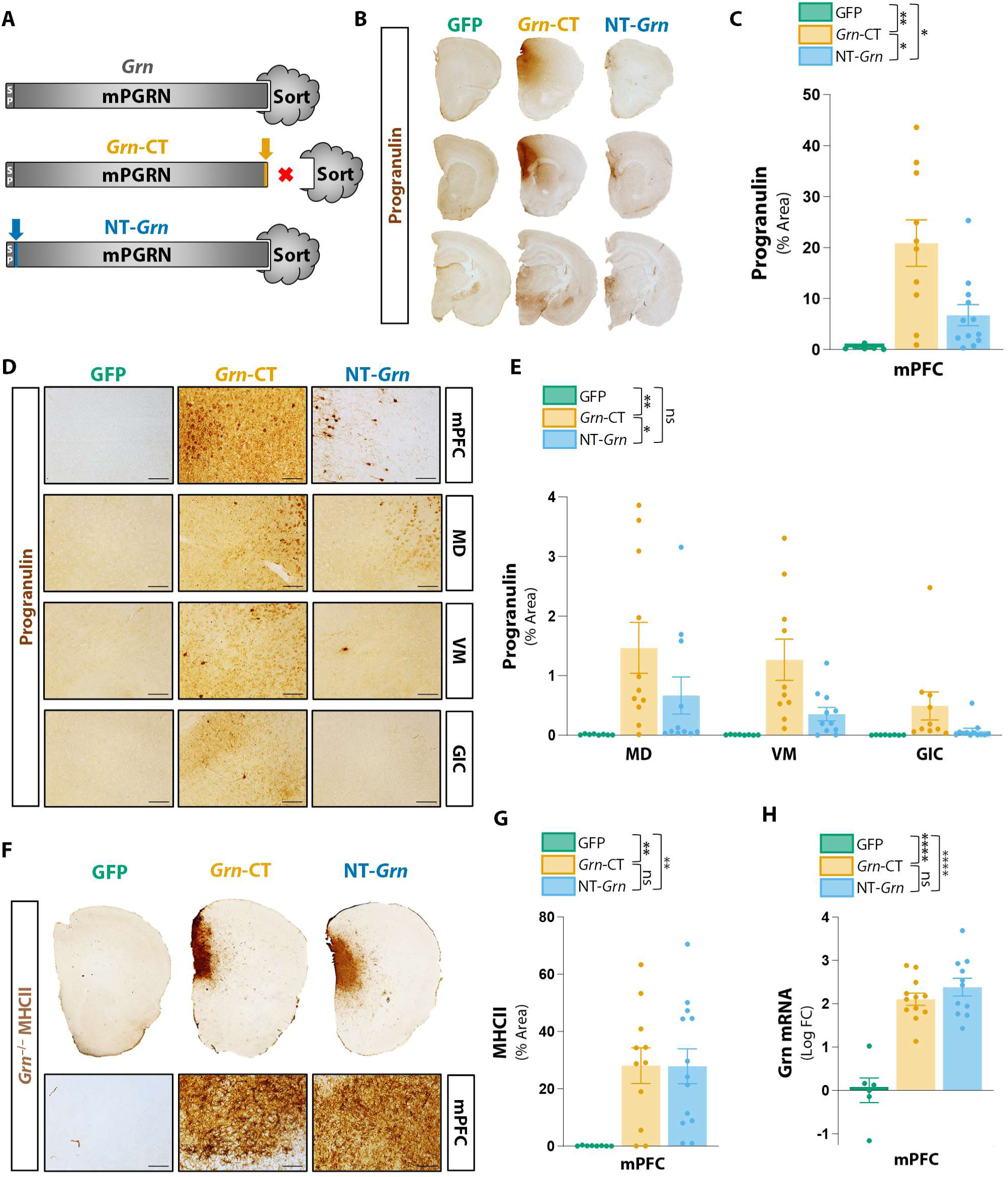
Blocking the progranulin C-terminus increases levels of AAV-derived progranulin. (**A**) *Grn*-CT has a tag at the progranulin C-terminus that blocks sortilin binding, unlike NT-*Grn* which has a tag at the N-terminus. (**B to E**) Progranulin immunohistochemistry in *Grn*^−/−^ mice treated with AAVs in the medial prefrontal cortex (mPFC) at 8–12 months of age and euthanized 8–11 weeks later. (**B**) Representative low-power sections. (**C**) Quantification of mPFC progranulin showing higher levels in *Grn*-CT–treated mice. Brown-Forsythe ANOVA, *P* = 0.0014; Tamhane’s T2 MC test: GFP vs. *Grn*-CT (*P* = 0.0045), GFP vs. NT-*Grn* (*P* = 0.0321), *Grn*-CT vs. NT-*Grn* (*P* = 0.0430); *n* = 6–12 mice per group. (**D**) Representative 20X micrographs of mPFC, mediodorsal (MD) and ventromedial (VM) nucleus of thalamus, and granular insular cortex (GIC). (**E**) Quantification showing higher progranulin in *Grn*-CT-treated mice. Restricted mixed-effects model, (*P* = 0.0045 for region effect, *P* = 0.0027 for AAV effect); Tukey’s MC test: GFP vs. *Grn*-CT (*P* = 0.0027), GFP vs. NT-*Grn* (*P* = 0.4175), *Grn*-CT vs. NT-*Grn* (*P* = 0.0315); *n* = 7–12 mice per group. (**F and G**) MHCII immunohistochemistry. (**F**) Representative low-power and 20X micrographs. (**G**) Quantification showing increased mPFC MHCII in both *Grn*-CT– and NT-*Grn*–treated mice. Brown-Forsythe ANOVA, *P* = 0.0045; Tamhane’s T2 MC test: GFP vs. *Grn*-CT (*P* = 0.0036), GFP vs. NT-*Grn* (*P* = 0.0020), *Grn*-CT vs. NT-*Grn* (*P* > 0.9999); *n* = 7–13 mice per group. (**H**) *Grn*-CT and NT-*Grn* similarly increased *Grn* mRNA. One-way ANOVA, *P* = 0.0008; Tukey’s MC test: GFP vs. *Grn*-CT (*P* < 0.0001), GFP vs. NT-*Grn* (*P* < 0.0001), *Grn*-CT vs. NT-*Grn* (*P* = 0.5273); *n* = 6–12 mice per group. Scale bars, 100µm.

To determine the effects of C-terminal blockade on levels of AAV-derived progranulin, we measured progranulin by immunohistochemistry 8–10 weeks after treatment. Consistent with our previous observations, *Grn*-CT robustly increased progranulin in the mPFC (Fig. 1B to D) and in other regions more distant from the injection site (Fig. 1B, D, and E). In contrast, NT-*Grn*–treated mice had significantly lower levels of AAV-derived progranulin than *Grn*-CT– treated mice, both in the mPFC (Fig. 1B and C) and in regions distant from the mPFC, including the mediodorsal nucleus of the thalamus (MD), ventromedial nucleus of the thalamus (VM), and granular insular cortex (GIC) (Fig. 1D and E). Qualitatively, *Grn*-CT yielded higher progranulin staining in the interstitial space than NT-*Grn*, which could be consistent with the predicted effects of C-terminal blockade reducing cellular uptake and degradation (Fig. 1D).

We considered several potential reasons for the difference in progranulin levels between *Grn*-CT and NT-*Grn*. In prior work, administering *Grn*-CT in *Grn^−/−^* mice produced a non-self immune reaction that curtailed the resulting progranulin levels (*20*). Thus, one potential explanation for the differences in AAV-derived progranulin levels between *Grn*-CT and NT-*Grn* could be higher immunogenicity of NT-*Grn* relative to *Grn*-CT, resulting in decreased NT-*Grn* derived progranulin. To assess immunogenicity of the AAV-*Grn* vectors we examined immunohistochemistry for major histocompatibility complex class II (MHCII), a marker of antigen presentation, at the injection site (Fig. 1F). There was no difference between *Grn*-CT– and NT-*Grn*–induced MHCII upregulation (Fig. 1G). Furthermore, neither *Grn*-CT nor NT-*Grn* had any measurable effects on body weight, brain weight, or survival relative to GFP-treated controls (fig. S2). These data provide no support for the idea of differential immunogenicity or toxicity between *Grn*-CT and NT-*Grn*.

Another potential explanation for the higher progranulin levels produced by *Grn*-CT would be if the *Grn*-CT vector had higher transduction efficiency and produced more *Grn* transcript than the NT-*Grn* vector, despite injection of equal viral genomes. To address this, we measured *Grn* mRNA levels by quantitative real-time PCR. *Grn* mRNA levels were equivalent between mice treated with *Grn*-CT and NT-*Grn* (Fig. 1H). Thus, for a given level of *Grn* mRNA produced, *Grn*-CT yielded higher progranulin protein levels than NT-*Grn*. The most likely explanation for the higher levels resulting from *Grn*-CT is the blockade of sortilin binding resulting in less progranulin degradation.

### Blocking the progranulin C-terminus does not enhance reduction of neuronal lipofuscinosis

Since blocking the progranulin C-terminus increased levels of AAV-derived progranulin both at and distant from the injection site, we asked whether it also enhanced the ability of AAV-progranulin to reduce pathology. Because they are post-mitotic, neurons are particularly susceptible to age-related lipofuscin accumulation (*30*). Like patients with FTD-*GRN*, progranulin-deficient mice exhibit age-dependent neuronal lipofuscinosis (*31–33*) detectable by autofluorescence or immunostaining for subunit C of mitochondrial ATP synthase (SCMAS). Previous work demonstrated that *Grn*-CT injection in the mPFC reduces lipofuscinosis in the ventral posterolateral and ventral posteromedial (VPM/VPL) nuclei of the thalamus and the CA3 pyramidal cell layer of the hippocampus (*20*).

To determine if blocking the progranulin C-terminus affects AAV-progranulin–mediated correction of neuronal lipofuscinosis, we first measured autofluorescence in these areas as well as layers II/III of the neocortex. Interestingly, both *Grn*-CT and NT-*Grn* ameliorated autofluorescence to a similar extent relative to GFP-treated controls (Fig. 2A and B). For an additional measure of neuronal lipofuscinosis we also immunostained for SCMAS, which more specifically labels neuronal lipofuscin in most forms of neuronal ceroid lipofuscinosis (*34*). As previously described (*20*), treatment with *Grn*-CT decreased SCMAS pathology relative to GFP. Consistent with the autofluorescence, the effects on NT-*Grn* on SCMAS pathology were similar to those of *Grn*-CT (Fig. 2C and D). Together, these data indicate that AAV-progranulin does not require C-terminal modification to reduce neuronal lipofuscinosis in this model of global progranulin insufficiency.

**Fig. 2.**
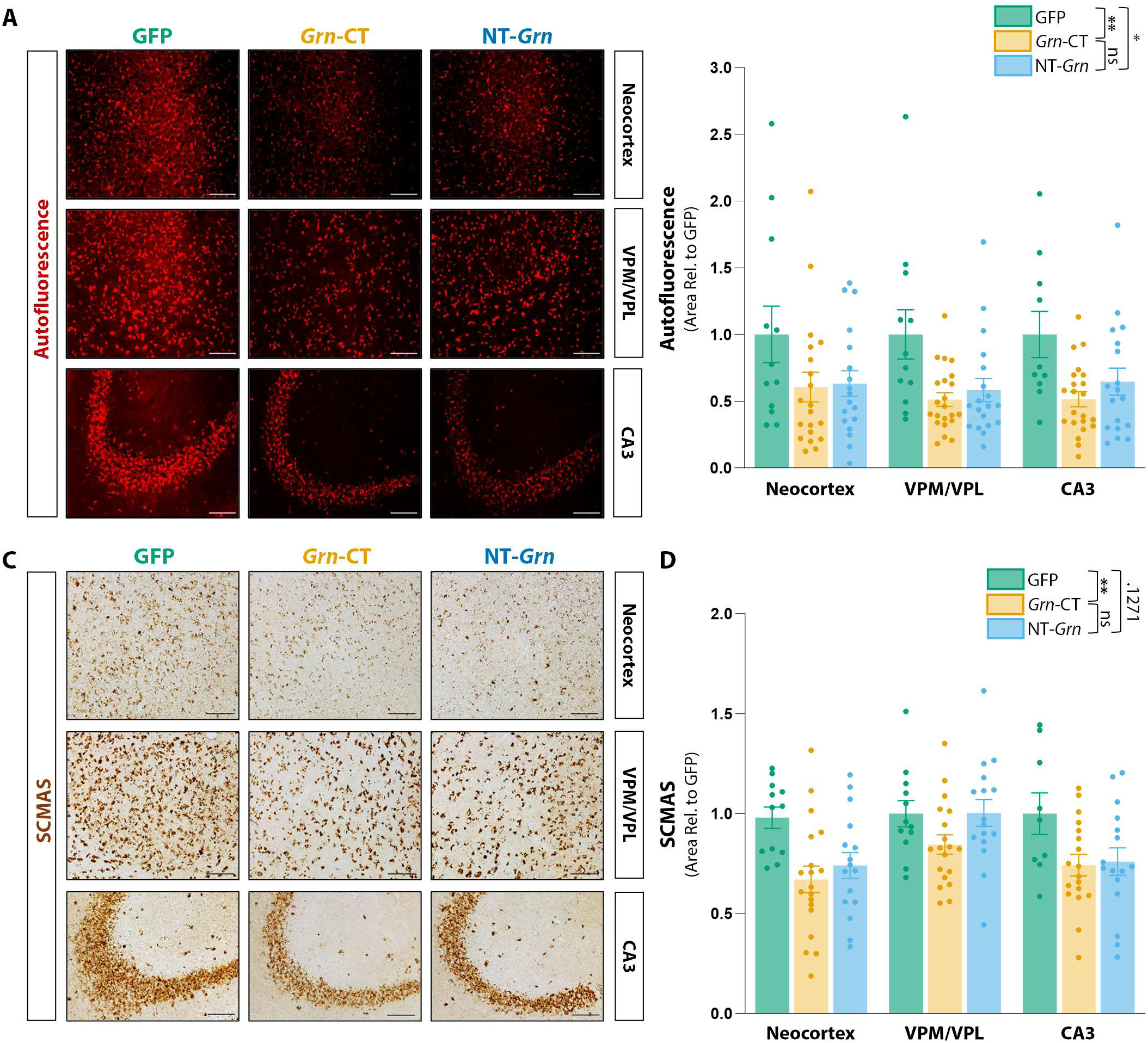
Both *Grn*-CT and NT-*Grn* reduce neuronal lipofuscinosis. **(A)** Representative 20X images of neuronal autofluorescence in Layer II of neocortex, ventral posteromedial and ventral posterolateral nucleus of thalamus (VPM/VPL), and hippocampal area CA3 in brains of *Grn*^−/−^ mice treated with GFP, *Grn*-CT and NT-*Grn*. **(B)** Both *Grn-*CT and NT-*Grn* treatment reduced autofluorescent lipofuscin pathology in neuron-heavy regions of the brain. Restricted maximum likelihood (REML) mixed-effects model (*P* = 0.8259 for region effect, *P* = 0.0064 for AAV effect); Tukey’s MC test: GFP vs. *Grn*-CT (*P* = 0.0055), GFP vs. NT-*Grn* (*P* = 0.0333), *Grn-*CT vs. NT-*Grn* (*P* = 0.7742); *n* = 13–22 mice). **(C)** Representative 20X micrographs of SCMAS immunohistochemistry in Layer II of the neocortex, VPM/VPL, and CA3. **(D)** Both *Grn*-CT and NT-*Grn* treatment reduced SCMAS pathology in neuron-heavy regions of the brain. Restricted maximum likelihood (REML) mixed-effects model (*P* = 0.0002 for region effect, *P* = 0.0116 for AAV effect); Tukey’s MC test: GFP vs. *Grn*-CT (*P* = 0.0083), GFP vs. NT-*Grn* (*P* = 0.1271), *Grn-* CT vs. NT-*Grn* (*P* = 0.4638); *n*=12–19 mice. Scale bars, 100µm.

### Blocking the progranulin C-terminus enhances reduction of microglial pathology

In addition to lipofuscinosis, *Grn*^−/−^ mice also exhibit age-dependent microgliosis. We previously demonstrated that delivery of *Grn*-CT to the mPFC of *Grn*^−/−^ mice rescues CD68 pathology in regions distant from the injection site, including motor cortex, CA3, and VPM/VPL of the thalamus (*20*). To determine if blocking the progranulin C-terminus is critical for this effect, we quantified CD68 immunoreactivity in these and additional brain regions of *Grn*^−/−^ mice treated with GFP, *Grn*-CT, or NT-*Grn*. Across these regions, *Grn*-CT, but not NT-*Grn*, reduced CD68 pathology (Fig. 3, A and B, fig. S3A).

**Fig. 3.**
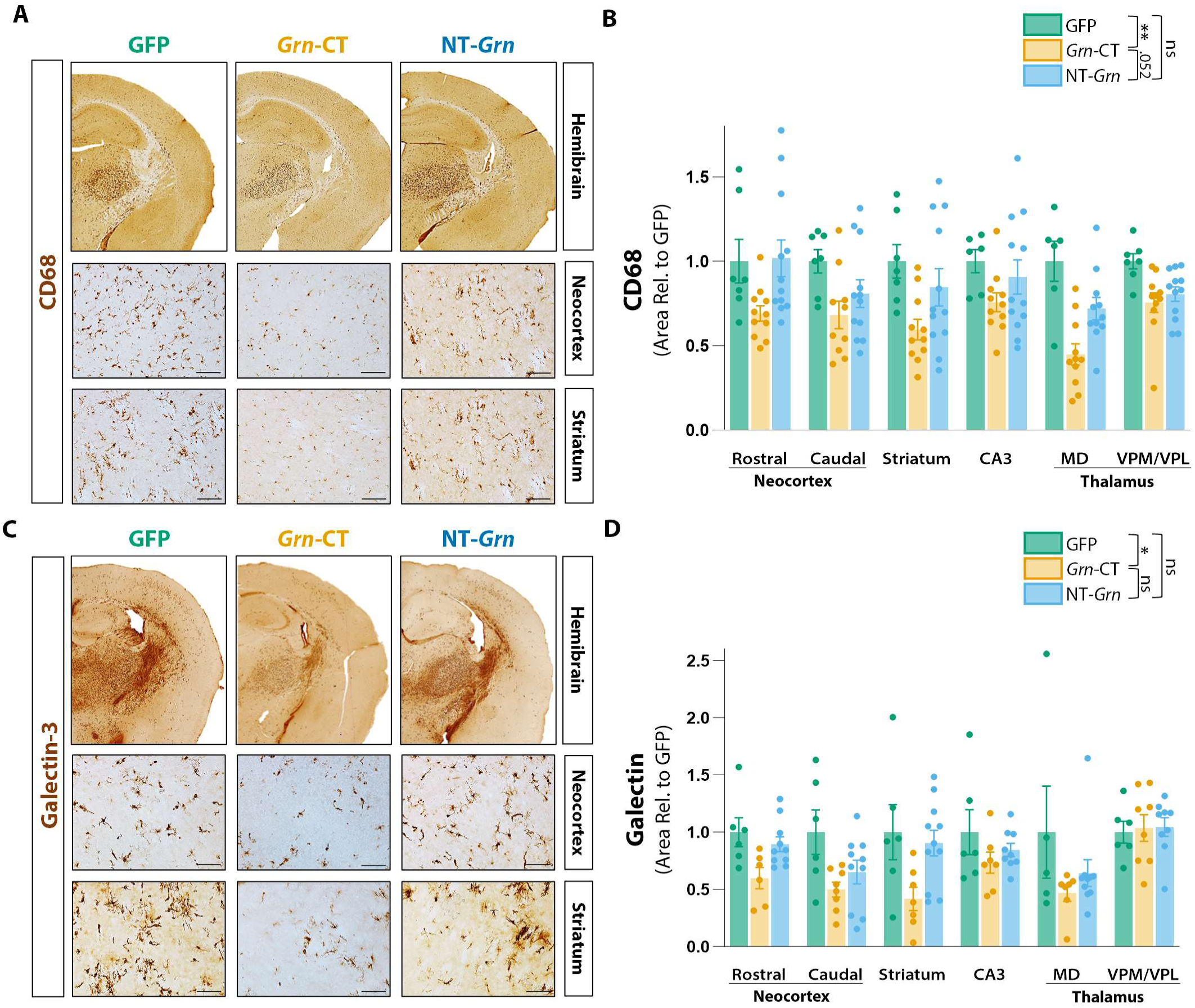
Blocking the progranulin C-terminus enhances AAV-Progranulin mediated reduction of neuroinflammation. **(A)** Representative hemibrain and 20X micrographs of cortical, hippocampal, thalamic, and striatal CD68 pathology in GFP, *Grn*-CT, and NT-*Grn* treated mice. **(B)** *Grn*-CT, but not NT-*Grn,* reduced CD68 pathology relative to GFP in areas of the neocortex, CA3, striatum, and thalamus. Restricted maximum likelihood (REML) mixed-effects model (*P* = 0.0223 for region effect, *P* = 0.0030 for AAV effect); Tukey’s MC test: GFP vs. *Grn*-CT (*P* = 0.0027), GFP vs. NT-*Grn* (*P* = 0.2600), *Grn*-CT vs. NT-*Grn* (*P* = 0.0521); *n*= 7–12 mice. **(C)** Representative hemibrain and 20X micrographs of cortical, hippocampal, thalamic, and striatal Galectin pathology in GFP, *Grn*-CT, and NT-*Grn* treated mice. **(D)** *Grn-*CT, but not NT*-Grn,* reduced Galectin pathology relative to GFP in the neocortex, striatum, CA3, and medial dorsal (MD) and VPM/VPL thalamus. Restricted maximum likelihood (REML) mixed-effects model (*P* = 0.0036 region effect, *P* = 0.0306 for AAV effect); Tukey’s MC test: GFP vs. *Grn*-CT (*P* = 0.0251), GFP vs. NT-*Grn* (*P* = 0.3786), *Grn*-CT vs. NT-*Grn* (*P* = 0.1948); *n*= 6–12 mice. Scale bars, 100µm.

To further investigate AAV-*Grn–*mediated rescue of microgliosis, we immunostained for Galectin-3, a β-galactoside–binding lectin found in reactive myeloid lysosomes with neuroinflammation (*35, 36*). Galectin-3 is upregulated by microglia in brains of aged *Grn*^−/−^ mice and is increased in brains of patients with FTD-*GRN* (*37*). As with CD68, treatment with *Grn*-CT, but not with NT-*Grn*, reduced cortical, striatal, thalamic, and hippocampal Galectin-3 pathology relative to GFP (Fig. 3, C and D, fig. S3B). Together, these data indicate that blocking the progranulin C-terminus enhances AAV-*Grn*–mediated reduction of neuroinflammation caused by progranulin deficiency.

### Blocking the progranulin C-terminus enhances reduction of microglial lipofuscinosis

While both NT-*Grn* and *Grn*-CT reduced neuronal lipofuscinosis (Fig. 2), the fact that only *Grn*-CT had significant effects on microgliosis caused by progranulin deficiency (Fig. 3) prompted us to look specifically at lipofuscinosis in microglia. To measure microglial lipofuscinosis we examined autofluorescence within Iba1-defined regions of interest in confocal images. To prevent neuronal lipofuscin from confounding measurements of microglial lipofuscinosis, we imaged regions with high microglia-to-neuron ratios, including layer I of the motor cortex and the stratum radiatum/stratum lacunosum-moleculare (SR/SLM) layers of hippocampal area CA3. In Layer I of the motor cortex, *Grn*-CT, but not NT-*Grn*, reduced microglial lipofuscinosis relative to GFP (Fig. 4, A and B). Similarly in the more distant region of hippocampal area CA3, *Grn*-CT but not NT-*Grn* treatment reduced microglial lipofuscinosis relative to GFP (Fig. 4, C and D). Together these data indicate that blocking the progranulin C-terminus is important for the ability of AAV-progranulin to ameliorate lysosomal pathology in microglia both at and distant from the injection site.

**Fig. 4.**
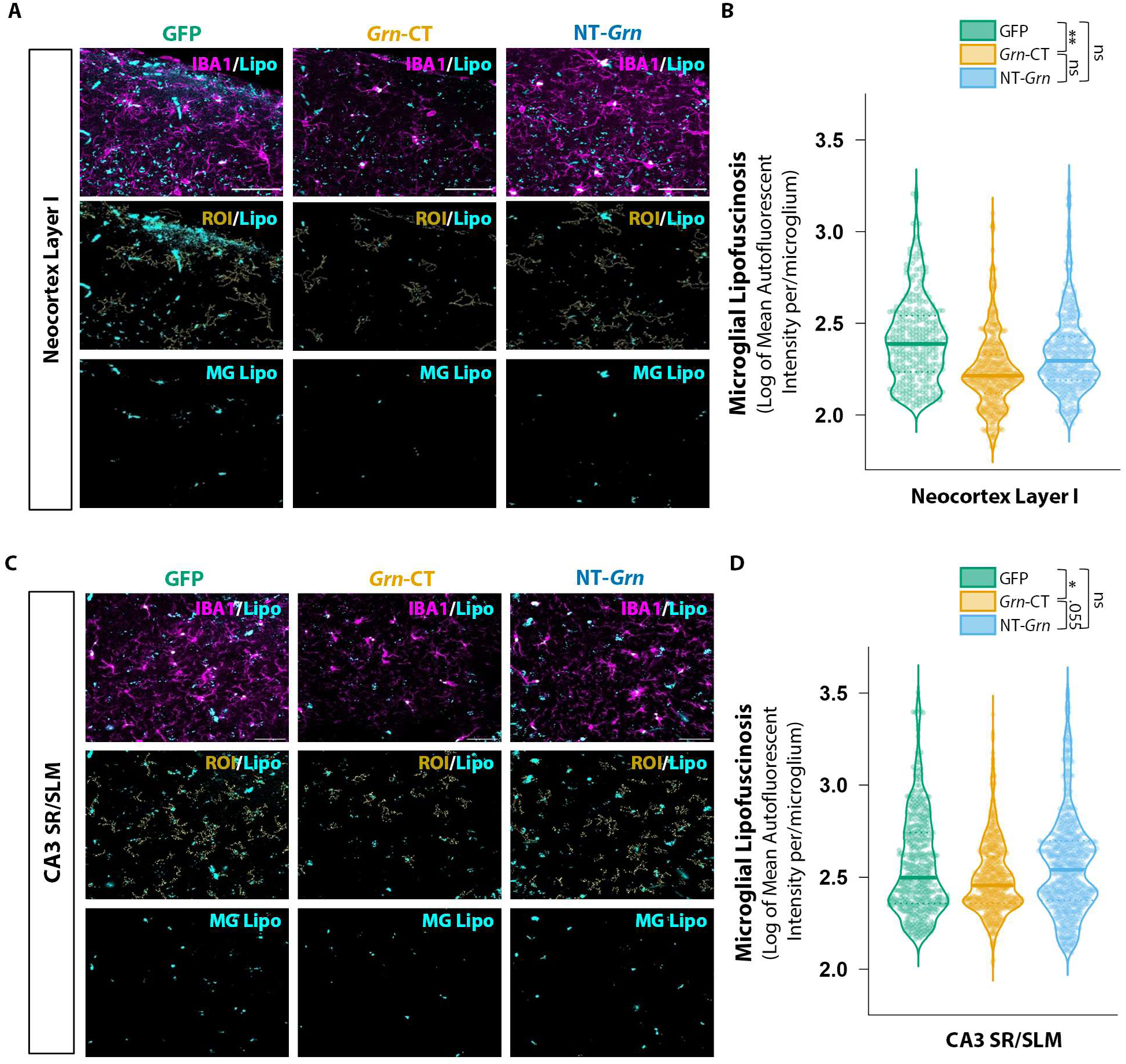
Blocking the progranulin C-terminus enhances AAV-Progranulin mediated reduction of microglial lipofuscinosis. **(A)** Representative images of microglial lipofuscinosis in neocortex (M1) Layer I. **(B)** *Grn-*CT, but not NT-*Grn*, reduced microglial lipofuscinosis in neocortex relative to GFP. Linear-mixed effects model fit by maximum likelihood with mouse as a random effect; GFP vs. *Grn-*CT (*P* = 0.00321), GFP vs. NT-*Grn* (*P* = 0.1281), *Grn-*CT vs. NT-*Grn* (*P* = 0.1051); *n* = 10–15 mice. **(C)** Representative images of microglial lipofuscinosis in SR/SLM layer of CA3. **(D)** *Grn*-CT, but not NT-*Grn,* reduced microglial lipofuscinosis in the SR/SLM layer of CA3 relative to GFP. Linear-mixed effects model fit by maximum likelihood with mouse as a random effect; GFP vs. *Grn*-CT (*P* = 0.0262), GFP vs. NT-*Grn*, (*P* = 0.5957), *Grn*-CT vs. NT-*Grn* (*P* = 0.0549); *n* = 9–13 mice. Scale bars, 100µm.

### Blocking the progranulin C-terminus enhances correction of lipid abnormalities

Recent lipidomic studies highlight altered lipid composition, particularly the accumulation of gangliosides and loss of bis(monoacylglycero)phosphate (BMP) lipids in FTD-*GRN* patients and *Grn*^−/−^ mice (*21, 38, 39*). To determine if AAV-progranulin gene therapy can correct these abnormalities, we performed lipidomic analysis across multiple brain regions (neocortex, thalamus, and cerebellum) of *Grn*^−/−^ mice treated with AAV expressing GFP, *Grn*-CT, or NT-*Grn*, along with *Grn*^+/+^ controls treated with AAV-GFP.

Consistent with previous observations (*21, 38*), *Grn*^−/−^ mice had loss of BMP species and accumulation of gangliosides relative to *Grn*^+/+^ mice (Fig. 5, A–D). These changes were most apparent in the neocortex (Fig. 5A), where both BMPs and gangliosides are most abundant, but thalamus and cerebellum had similar changes (Fig. 5, B and C). Both *Grn*-CT and NT-*Grn* improved these abnormalities across all three regions, increasing BMP levels and promoting clearance of ganglioside accumulation relative to GFP-treated *Grn*^−/−^ mice (Fig. 5D).

**Fig. 5.**
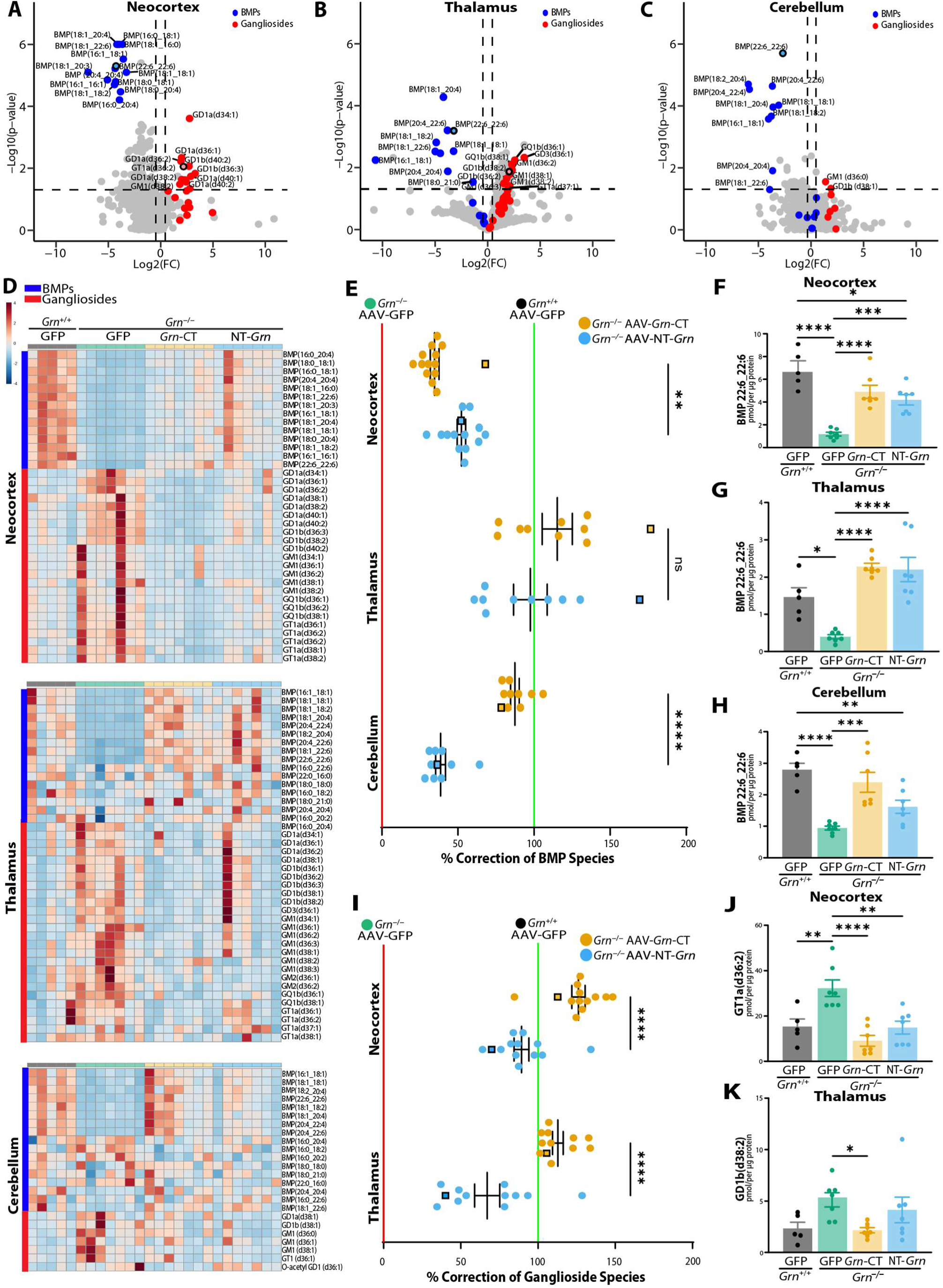
Blocking the progranulin C-terminus enhances correction of lipid abnormalities. (**A-C**) Volcano plots showing differential lipid levels in *Grn*^−/−^ vs *Grn*^+/+^ mice, with loss of BMP (blue) and ganglioside accumulation (red). (**D**) Heatmaps showing changes in the abundant species of BMPs (blue) and gangliosides (red). (**E**) Percent correction for all BMP species that were significantly different and ±1.5 log2-fold change in *Grn*^−/−^ vs. *Grn*^+/+^ mice (0% correction = level in GFP-treated *Grn*^−/−^ mice and 100% correction = level in GFP-treated *Grn*^+/+^ mice). Squares represent the lipids shown in F-H. Brown-Forsythe ANOVA, *P* < 0.0001; Tamhane T2 MC test: *Grn*-CT vs. *Grn*-NT Cortex (*P* = 0.0023); *Grn*-CT vs. *Grn*-NT Cerebellum (*P* < 0.0001). (**F–H**) Levels of the most abundant BMP species, di-22:6, in each region. (**F**) Neocortex. One-way ANOVA (*P* <0.0001); Tukey’s MC test: *Grn*^+/+^ GFP vs. *Grn*^−/−^ GFP (*P* < 0.0001); *Grn*^−/−^ GFP vs. *Grn*-CT (*P* < 0.0001); *Grn*^−/−^ GFP vs. *Grn*-NT (*P* = 0.0008). *Grn*-CT vs. *Grn*-NT (*P* = 0.7141). (**G**) Thalamus. One-way ANOVA (P <0.0001); Tukey’s MC test: *Grn*^+/+^ GFP vs. *Grn*^−/−^ GFP (*P* = 0.0117); *Grn*^−/−^ GFP vs. *Grn*-CT (*P* < 0.0001); *Grn*^−/−^ GFP vs. *Grn*-NT (*P* < 0.0001). *Grn*-CT vs. *Grn*-NT (*P* = 0.9927). (**H**) Cerebellum. One-way ANOVA (*P* <0.0001); Tukey’s MC test: *Grn*^+/+^ GFP vs. *Grn*^−/−^ GFP (*P* < 0.0001); *Grn*^−/−^ GFP vs. *Grn*-CT (*P* = 0.0005); *Grn*^−/−^ GFP vs. *Grn*-NT (*P* = 0.1467). *Grn*-CT vs. *Grn*-NT (*P* = 0.0770). (**I)** Percent correction for all gangliosides that were significantly different and ±1.5 log2-fold change in *Grn*^−/−^ vs *Grn*^+/+^ mice. Squares indicates most abundant gangliosides, shown in J,K. One-way ANOVA, *P* < 0.0001; Tukey’s MC test: *Grn*-CT vs. *Grn*-NT Cortex (*P* < 0.0001); *Grn*-CT vs. *Grn*-NT Thalamus (*P* < 0.0001). (**J,K**) Levels of the most abundant gangliosides in each region. (**J**) GT1a (d36:2) in neocortex. One-way ANOVA (*P* = 0.0001;); Tukey’s MC test: *Grn*^+/+^ GFP vs. *Grn*^−/−^ GFP (*P* = 0.0069); *Grn*^−/−^ GFP vs. *Grn*-CT (*P* < 0.0001); *Grn*^−/−^ GFP vs. *Grn*-NT (*P* = 0.0023). *Grn*-CT vs. *Grn*-NT (*P* = 0.5220). (**K**) GD1b(d38:2) in thalamus. One-way ANOVA (*P* = 0.0344;); Tukey’s MC test: *Grn*^+/+^ GFP vs. *Grn*^−/−^ GFP (*P* = 0.0983); *Grn*^−/−^ GFP vs. *Grn*-CT (*P* = 0.04520452); *Grn*^−/−^ GFP vs. *Grn*-NT (*P* = 0.7091). *Grn*-CT vs. *Grn*-NT (*P* = 0.3221). *n* = 5-7 mice per group for all comparisons.

Interestingly, *Grn*-CT overall produced a stronger beneficial effect. For BMP, both *Grn*-CT and NT-*Grn* boosted BMPs in neocortex and thalamus (Fig. 5, D and E). The two constructs had similar effects overall in the thalamus and on the most abundant BMP species, di-22:6 BMP, in both neocortex and thalamus (Fig. 5 F and G), although NT-*Grn* increased most BMP species slightly better than Grn-CT in the neocortex (Fig. 5E), driven by one mouse visible as a dark red vertical line in Fig. 5D. But in the cerebellum, the region most distant from the injection site, *Grn*-CT had a much stronger effect on correcting loss of BMP (Fig. 5, E and H), consistent with the finding that *Grn*-CT spreads further than NT-*Grn*.

For gangliosides, *Grn*-CT provided stronger reversal of the accumulation in both cortex and thalamus, both for gangliosides overall (Fig. 5I) and for the most abundant ganglioside in each region (Fig. 5, J and K). Too few gangliosides accumulated in cerebellum to perform this comparison.

### Blocking the progranulin C-terminus prevents age-dependent elevation of plasma NfL

Since blocking the progranulin C-terminus improved AAV-progranulin–mediated correction of neuroinflammation and lipid metabolism in several areas of the brain, we determined its effects on a biomarker of neurodegeneration. Plasma NfL measured on the Quanterix SIMOA platform is a highly sensitive and translationally relevant biomarker of neurodegeneration that, by disease progression modeling, is elevated up to ten years before symptom onset in carriers of *GRN* mutations (*40*).

To determine if blocking the progranulin C-terminus improved AAV-progranulin– mediated reduction of plasma NfL, we collected plasma from GFP-treated *Grn^+/+^* mice and from GFP-treated, *Grn-*CT*–*treated, and NT-*Grn–*treated *Grn^−/−^* mice. Relative to GFP-treated *Grn^+/+^* mice, GFP-treated *Grn^−/−^* mice had elevated plasma NfL indicative of neurodegeneration (Fig. 6). However, while *Grn*-CT reduced plasma NfL relative to GFP, NT-*Grn* did not (Fig. 6), suggesting that blocking the progranulin C-terminus is critical for the ability of AAV-progranulin gene therapy to prevent the progression of neurodegeneration in this mouse model of FTD-*GRN*.

**Fig. 6.**
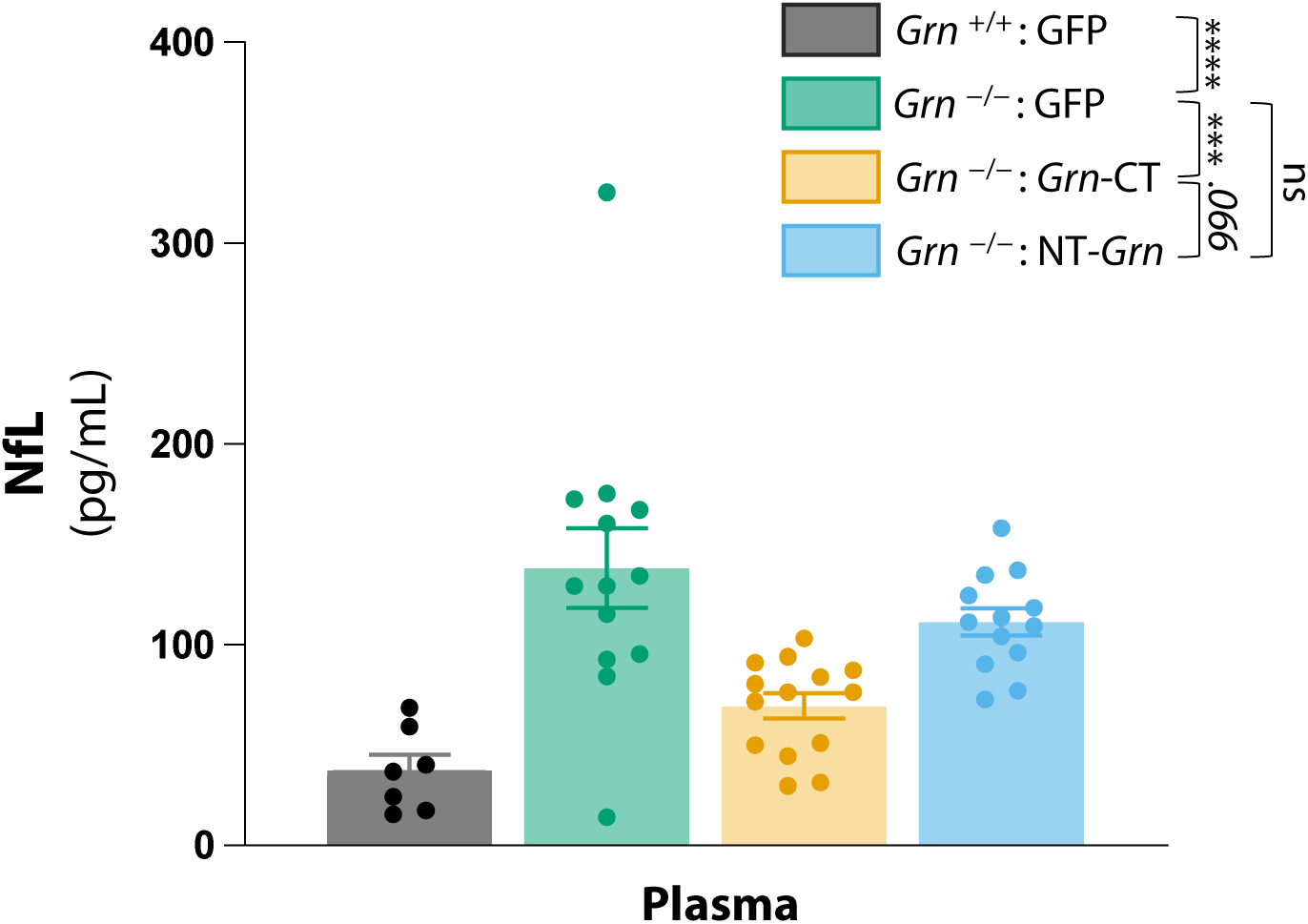
Blocking the progranulin C-terminus enhances AAV-Progranulin mediated improvement of plasma NfL. *Grn*-CT, but not NT-*Grn,* reduced plasma NfL in *Grn^−/−^* mice. One-way ANOVA, *P* < 0.0001; Tukey’s MC test, *Grn^+/+^* GFP vs. *Grn^−/−^* GFP (*P* < 0.0001); *Grn^+/+^* GFP vs *Grn^−/−^ Grn*-CT (*P* = 0.3821); *Grn^+/+^* GFP vs *Grn^−/−^* NT-*Grn* (*P* = 0.0034); *Grn^−/−^* GFP vs. *Grn^−/−^ Grn*-CT *(P* = 0.0008), *Grn^−/−^* GFP vs. *Grn^−/−^* NT-*Grn* (*P* = 0.3916), *Grn^−/−^ Grn*-CT vs. *Grn^−/−^* NT-*Grn* (*P* = 0.0666); *n* = 7–14.

## DISCUSSION

The goal of this study was to determine if blocking the progranulin C-terminus, which harbors the sortilin binding site, enhances the efficacy of AAV-progranulin gene therapy. At the same dose, AAV expressing progranulin with the C-terminus blocked yielded more progranulin protein both at the injection site and in more distant regions than AAV expressing unblocked progranulin (Fig. 1). Blocking the progranulin C-terminus was critical for the ability of AAV-progranulin to reduce microglial pathology, including multiple measures of microgliosis (Fig. 3) and microglial lipofuscinosis (Fig. 4). While unblocked progranulin improved both ganglioside accumulation and loss of BMP, blocking the progranulin C-terminus increased the effectiveness of correcting both lipid abnormalities further from the injection site (Fig. 5). Finally, only progranulin with the C-terminus blocked improved plasma NfL, a translationally relevant biomarker of neurodegeneration (Fig. 6). Altogether, these data indicate that blocking the progranulin C-terminus, which we previously showed prevents sortilin interactions in this vector (*20*), improves the effectiveness of progranulin gene therapy.

Our data add to growing evidence that sortilin binding has an important influence on progranulin therapies. At one level, it is perhaps surprising that it would be desirable to block progranulin’s ability to bind sortilin, a major neuronal receptor that traffics progranulin to the lysosome where it is active (*26*). But genetic ablation of sortilin boosts brain and peripheral progranulin in wild-type and progranulin heterozygous knockout (*Grn*^+/–^) mice without apparent adverse effects (*26*). Several different approaches targeting the sortilin-progranulin axis have shown potential. A small molecule that reduces sortilin raised extracellular progranulin levels in cell lines (*23*). Another small molecule that blocks sortilin-mediated progranulin uptake is in a phase 1 clinical trial (NCT06226064). Anti-sortilin antibodies, which decrease progranulin-sortilin interactions by reducing sortilin levels, increase progranulin levels in mouse models (*28, 29*) and FTD-*GRN* patients (*41*). A phase 3 clinical trial of one such anti-sortilin antibody, latozinemab, is underway for FTD-*GRN* (NCT04374136).

One might wonder if increasing total (or extracellular) progranulin levels by blocking sortilin interactions could be counterproductive if it prevented progranulin from reaching the lysosome, where it has important functions (*42*). But sortilin is not necessary for progranulin’s lysosomal localization and *Grn*-CT localizes to neuronal lysosomes despite not binding sortilin (*19, 20*). This is likely because other progranulin receptors can also traffic progranulin to the lysosome, including via mannose 6-phosphate receptor and its interactions with prosaposin (*43, 44*). In addition to the data shown here that blocking sortilin interactions in AAV-progranulin has beneficial effects in mouse models, the anti-sortilin antibody latozinemab corrected social deficits in *Grn*^+/–^ mice (*29*). While no efficacy data in FTD-*GRN* patients are yet published for latozinemab, there were no safety signals in phase 1 or 2 to preclude its advance into the currently active phase 3 trial. Thus, the available data suggest, perhaps counterintuitively, that blocking progranulin binding to one of its neuronal receptors, sortilin, is beneficial even in a condition caused by progranulin loss of function.

The effects of blocking the progranulin C-terminus differed between outcome measures. With some measures, AAV-progranulin had beneficial effects regardless of whether the C-terminus was blocked. This was the case for correction of neuronal lipofuscinosis (Fig. 2) and may indicate that neurons are sensitive to even very low levels of progranulin, which are enough to correct these abnormalities. Consistent with this hypothesis, conditional deletion of progranulin in both excitatory neurons and microglia, which did not completely eliminate brain progranulin, did not cause neuronal lipofuscinosis (*45*). On the other hand, microglia-related outcome measures were much more sensitive to the effects of C-terminal blockade (Figs 3-4). This is likely related to the fact that microglia, unlike neurons, do not use sortilin or prosaposin for lysosomal localization of progranulin (*46*). Thus, blocking sortilin-mediated uptake by neurons via C-terminal blockade would leave more progranulin available to cross-correct microglia.

The observation that *Grn*-CT but not NT-*Grn* corrected the increase in plasma NfL is interesting for several reasons. First, increased NfL is a translational biomarker of neurodegeneration that the US Food and Drug Administration (FDA) recently used as the basis for an accelerated approval decision (*47*). The FDA has accepted a letter of intent to qualify NfL as a clinical trial biomarker for FTD. Second, as a plasma measure it reflects an integrated readout of effects throughout the brain, unlike other neuropathological measures that must be examined in specific brain regions. And third, the dissociation between effects on neuronal lipofuscinosis, which was equally corrected by both *Grn*-CT and NT-*Grn* (Fig. 2), and NfL, which was corrected only by *Grn*-CT (Fig. 6), suggests that neuronal lipofuscinosis is not a major driver of neurodegeneration in *Grn^−/−^* mice. The fact that the effects of *Grn*-CT and NT-*Grn* on NfL correlated better with their effects on microglial pathology (Figs. 3 and 4) are consistent with the hypothesis that microglial abnormalities contribute to neurodegeneration in FTD-*GRN*.

A key consideration for translation of AAV-progranulin gene therapy from mouse models to FTD-*GRN* patients is ensuring adequate spread of progranulin throughout the much larger human brain. Despite the “frontotemporal” in its name, FTD affects networks distributed throughout the brain, including the cerebellum (*48, 49*). AAV serotype and route of administration have traditionally been the primary variables for modulating distribution of gene therapies throughout the brain. Our study demonstrates that AAV cargo modifications can also have important effects on distribution, with C-terminal blockade increasing levels and spread of progranulin (Fig. 1).

Our data suggest that the features of two major therapeutic approaches for FTD-*GRN*, progranulin gene therapy and anti-sortilin therapy, can be combined using gene therapy to deliver C-terminally modified progranulin that lacks sortilin binding. In mouse models, this approach using C-terminal blockade facilitated broader spread of progranulin throughout the brain, reduced a biomarker of neurodegeneration, and enabled cross-correction of microglial abnormalities. Three companies have entered clinical trials of progranulin gene therapy, all with unmodified C-terminus (NCT04408625, NCT04747431, and NCT06064890). One of these studies recently reported interim results, effectively raising cerebrospinal fluid progranulin, but was not powered to determine efficacy (*50*). Our findings suggest that the effectiveness of next-generation progranulin gene therapy approaches may be maximized by delivering a cargo with C-terminal modifications that prevent sortilin interactions.

## MATERIALS AND METHODS

### Study Design

This was a controlled laboratory experiment to determine if AAV gene therapy delivering progranulin with the C-terminus blocked vs. unblocked had different effects in mouse models of progranulin deficiency. We delivered AAV expressing GFP control, C-terminally blocked progranulin (*Grn*-CT), or progranulin with an N-terminal tag and unblocked C-terminus (NT-*Grn*) via bilateral intraparenchymal injection into the prefrontal cortex of *Grn*^−/−^ mice. Outcome measures included progranulin levels and spread, lipofuscinosis, microgliosis, gangliosidosis measured by lipidomics, and neurodegeneration measured by plasma NfL biomarkers. Sample size was determined by power analyses using the *pwr* package in R, based on effect sizes and variability observed in our previously published related work (*19, 20*). The unit of investigation was individual mice except for microglial lipofuscinosis where it was individual microglial cells fit into a linear model with mouse as a random effect. Altogether for the various outcome measures, we analyzed five separate cohorts of mice, each with a sample size of 22–36 mice (7–13 in each of the three groups), for a total of 139 mice (37–49 in each of the three groups). Four mice in each group died between AAV injection and tissue collection 8–10 weeks later, so the final sample size analyzed postmortem was 127 mice (33–45 in each of the three groups).

No data or outliers were excluded, but in a few isolated cases data for individual mice was missing due to technical issues with that sample. The final *n* for each outcome measure is reported in figure legends and the number of technical replicates for each experiment is reported in the methods section. One cohort used for lipidomic and NfL biomarker analyses also included *Grn*^+/+^ mice treated with AAV-GFP control. Several outcome measures were internally replicated in separate cohorts. Randomization to treatment group was stratified by sex and age to achieve balanced allocation of these characteristics between groups. Investigators and animal care staff were blinded to genotype/treatment during data collection and analysis. Many analyses utilized semi-automated or completely automated workflows as detailed below. Order of testing/analysis was stratified by group to eliminate confounding by time-of-day or day-of-testing.

### Animals

Progranulin knockout mice on a congenic C57BL/6J background were generated as described previously (*51*). Cohorts of *Grn*^−/−^ mice used for this study were obtained from crossing *Grn*^+/–^ x *Grn*^+/–^ mice or *Grn*^−/−^ x *Grn*^−/−^ mice. The cohort that included *Grn*^+/+^ mice was obtained from crossing *Grn*^+/–^ x *Grn*^+/–^ mice. Each cohort included both male and female mice. All mice were maintained on a 12h:12h light/dark cycle and given *ad libitum* access to food and water in a barrier facility accredited by the Association for Assessment and Accreditation of Laboratory Animal Care. Mice were aged to 8–12 months before stereotaxic injection with AAV vectors and sacrificed 8–10 weeks after injection (see below). All experiments were approved by the Institutional Animal Care and Use Committee of the University of Alabama at Birmingham.

### Generation of AAVs

Both *Grn*-CT and NT-*Grn* were generated by cloning the RNA-coding sequence of the mouse progranulin gene (*Grn*) with a c-terminal *myc* tag (in the case of *Grn*-CT) or an N-terminal HA tag (in the case of NT-*Grn*) into a CIGW vector backbone, generously provided by Dr. David Standaert and Dr. Talene Yacoubian at the University of Alabama at Birmingham. For *Grn*-CT, the C-terminal tag was inserted after L593, the C-terminus of progranulin (RRID:Addgene_178944). For NT-*Grn*, the N-terminal tag was inserted after the signal peptide of progranulin. The CIGW backbone consists of CBA promoter, WPRE and rBG elements, and AAV2 inverted terminal repeats. Both *Grn*-CT (rAAV2-CBA-m*Grn*-Myc-WPRE-rBG) and NT-*Grn* (rAAV2-CBA-HA-m*Grn*-WPRE-rBG) were packaged into AAV1 vectors by the University of Pennsylvania Vector Core as described previously (*19*). The control AAV-GFP construct was an AAV1 construct with a CB7 promoter, WPRE, and rBG elements, purchased from the University of Pennsylvania Vector Core (AAV-GFP, #AV-1-PV1963, AAV1-CB7-CI-eGFP-WPRE-rBG).

### Stereotaxic delivery of AAVs

8-to 12-month-old mice were anesthetized with 5% isoflurane and then prepared for bilateral injection of AAV expressing *Grn*-CT, NT-*Grn*, or GFP via stereotaxic surgery as described previously (*19*). Briefly, the mouse head was shaved before its placement into a stereotaxic apparatus connected to a vaporizer for continuous delivery of isoflurane throughout the surgery. Mice were maintained on 1–2% isoflurane for the duration of the procedure. After anesthetization and before initial incision, mice received ocular ointment application and subcutaneous injection of 0.9% sterile saline, buprenorphine, and carprofen. After initial incision to expose the skull, a digital coordinate system and drill were used to create a burr hole at the location of the mPFC (coordinates +1.9 mm anterior and ± 0.3 mm lateral from bregma, −2.2 mm from the surface of the skull). 1uL of AAV (7.36 × 10^11^ genomes/mL) was infused into each hemisphere with a syringe pump (Harvard Apparatus) at a flow rate of 0.5 µL/min. Viruses were allowed to diffuse into the parenchyma for five minutes before the injection needle (Hamilton) was slowly withdrawn. Drill holes were sealed with bone wax and the incision wound was closed by surgical staples or silk sutures. Mice were allowed to recover on a heating pad and monitored every week for the duration of treatment. Surgical procedures and pre-and post-operative care were approved by the Institutional Animal Care and Use Committee of the University of Alabama at Birmingham.

### Tissue collection and subdissection

Mice were sacrificed 8–12 weeks after AAV injection by anesthesia with intraperitoneal injection of pentobarbital (100 mg/kg, Fatal Plus; Vortech Pharmaceuticals) and subsequent diaphragmectomy. Whole blood was collected through cardiac puncture and placed into tubes containing EDTA and spun 10,000 x *g* at 4°C for 10 minutes. Plasma was then placed into Protein Lo-bind tubes at −80°C until use. Mice were transcardially perfused with 0.9% saline before brain collection. Brains were isolated and hemisected at the sagittal sinus for fixation of one hemisphere and freezing of the other. Hemibrains used for sectioning and immunohistochemistry experiments were drop-fixed in 4% PFA for 48 hours and then transferred to PBS until sectioning (see below). Hemibrains used for qRT-PCR were flash frozen in dry ice and stored at −80°C until RNA isolation (see below).

Flash frozen hemibrains were sliced coronally at 1.0mm intervals with a stainless-steel adult mouse brain matrix (Zivic Instruments). Medial prefrontal cortex was subdissected from slices between the olfactory bulbs and emergence of corpus callosum and flash frozen on dry ice in 1.7mL Eppendorf tubes until RNA isolation.

### RNA isolation

Samples were homogenized with TRIZOL under a fume hood, shaken with chloroform, incubated for 5 minutes, and centrifuged at 4°C for 15 minutes at full speed to separate into aqueous and organic layers. The aqueous layer was incubated with glycogen and cold isopropyl alcohol for 15 minutes and centrifuged at 4°C for 25 minutes at full speed. The resulting pellet was resuspended in 75% ethanol (prepared in RNase-free water) and centrifuged at 4°C for 5 minutes at 7500 x *g*. The final pellet was then air-dried and resuspended in 25 µL of RNase-free water for subsequent cDNA synthesis and qRT-PCR.

### qRT-PCR

RNA was incubated with DNase I (Sigma) for 10 minutes and then combined with iScript cDNA synthesis reagents (BioRad) according to kit directions. cDNA synthesis was performed in a thermal cycler (BioRad) according to kit specifications. cDNA samples were then incubated with primers for mouse *Grn* mRNA (Taqman; Mm01245914_g1) and mouse *Actb* mRNA (Taqman; Mm02619580_g1). qRT-PCR was performed with the QuantStudio 5 thermal cycler (AppliedBiosystems) according to Taqman assay protocol specifications. *Grn* mRNA was normalized to *Actb* mRNA via the delta-delta CT method. Two technical replicates were run for each sample.

### Diaminobenzidine (DAB) immunohistochemistry

Fixed hemibrains were cryoprotected in a solution of 30% sucrose in PBS for 14–16 hours and sectioned into 30 *µ*m sections on a sliding microtome (Leica). For quantitative assessments of progranulin and microglial pathology, sections were incubated overnight in primary antibody (see below) at 4°C. The following day sections were incubated in species-matched biotinylated secondary antibody (Vector Labs, 1:1000) at room temperature followed by 30-minute incubation with avidin-biotin complex (Vectastain Elite; Vector Laboratories). Immunostaining was visualized via development with a solution of diaminobenzidine (MP Biomedicals) in 12% hydrogen peroxide.

*Antibodies for diaminobenzidine immunohistochemistry:*

| Marker | Product | Concentration | Source and Immunogen |
| --- | --- | --- | --- |
| Progranulin | R and D Systems<br>Cat# AF2557<br>RRID: AB_2114504 | 1:100 | Sheep polyclonal to recombinant mouse progranulin (Thr18-Leu589) |
| MHCII | Invitrogen<br>Cat# 14-5321-82<br>RRID: AB_467561 | 1:100 | Rat monoclonal; immunogen peptide information is proprietary |
| SCMAS | Abcam<br>Cat# ab181243<br>RRID: AB_2935765 | 1:250 | Rabbit monoclonal to ATP synthase F(0) complex subunit C1, mitochondrial (Human) |
| CD68 | BioRad<br>Cat# MCA1957<br>RRID: AB_2244437 | 1:500 | Rat monoclonal to purified concanavalin A acceptor glycoprotein from mouse P815 cells; peptide AA sequence is proprietary |
| Galectin-3 | R and D Systems<br>Cat# AF1197<br>RRID: AB_2234687 | 1:100 | Goat polyclonal to *E.coli*-<br>derived recombinant mouse<br>Galectin-3 (Ala2-Ile264) |

### Epifluorescent and brightfield microscopy

Progranulin, MHCII, SCMAS, CD68, and Galectin-3 levels were analyzed by imaging immunostained sections with a light microscope (Nikon) and CCD camera (Nikon). Lipofuscinosis in neuron-heavy regions of the brain was measured by mounting unstained hemibrain sections with DAPI-containing mounting medium (ProLong Diamond) for visualization of autofluorescence with an epifluorescent microscope (Nikon). Lipofuscin was visualized in the TRITC channel to avoid fluorescent signal from GFP and captured with a CCD camera (Andor Technologies). Brain regions of interest were imaged at 20X in most cases. The striatum and rostral neocortex regions of Galectin-3–labeled sections were imaged at 10X to account for the patchy, geographical distribution of Galectin pathology. 1-2 images of each region were analyzed for each mouse.

### Immunofluorescence

Sections were blocked for one hour and then incubated overnight with Rabbit Anti-Mouse Iba1 (Wako Chemicals USA, ordered via Fisher Scientific Cat# NC9288364, RRIDs: AB_839504, AB_10206679, AB_2665520; 1:500), and then incubated in AlexaFluor 647-conjugated Anti-Rabbit secondary (Invitrogen, Cat# A-21244, RRID:AB_2535812; 1:500) for 1–2 hours at room temperature in the dark. Sections were then incubated in DAPI (Invitrogen; 1:200) for 15 minutes before mounting with 70% glycerol on ColorFrost slides.

### Confocal microscopy

To avoid overlap with autofluorescent signal from brain lipofuscin, which has a wide emission spectrum between 490 to 660 nm (*52, 53*), an Alexa Fluor 647 conjugated secondary antibody (peak emission at 671 nm when excited with a 640 nm laser) was used to label Iba1 positive cells. To avoid overlap with GFP signal in GFP-treated mice, the 561nm laser was used to measure autofluorescent lipofuscin puncta. We used Sequential scanning microscopy with two separate optical configurations for Cy5 and DAPI/TRITC to minimize channel spillover between the TRITC and Cy5 lasers. For Layer I Cortex 17.00 *µ*m stacks were taken with the 60X Oil objective (86 images per stack, taken every 0.2 *µ*m) and for CA3: SLM 8.00 *µ*m stacks were taken with the 40X Oil objective (41 images per stack, taken every 0.2*µ*ms). 1-2 Z-stacks were imaged for each region, per mouse.

### Image analysis

#### DAB Immunostaining and Neuronal Lipofuscinosis

All brightfield images and epifluorescent lipofuscin images (for neuronal lipofuscinosis) were converted into binary representations and analyzed with Fiji via an automated workflow. Density of immunostaining and lipofuscinosis was quantified by applying a uniform threshold and measuring area occupied by immunopositive pixels within the applied threshold. Full hemibrain images were obtained with a slide scanner (PathScan Enabler, Meyer Instruments).

#### Microglial Lipofuscinosis

Multichannel images of Iba1, DAPI and autofluorescent lipofuscin were analyzed via a semi-automated workflow, in which images were first split in Fiji to isolate DAPI, lipofuscin, and Iba1 channels. Iba1 stacks were then isolated and processed with the Fiji “Desplecking” function to despeckle all images within the stack and the Fiji “Remove Outliers” function with a radius of 5.0 µm and a threshold of 50 to remove bright pixel outliers. The processed Iba1 stacks were then compressed into a maximum intensity projection and thresholded by pixel value to isolate Iba1-positive staining. The “Analyze Particles” function with size exclusion of 15–900 pixels was used to segment Iba1-positive cells, and subsequent ROIs were visually inspected to minimize the inclusion of ROIs of multiple cells and ROIs of portions of cells without cellular somas. All ROIs were then saved into ROI sets for each individual mouse. Lipofuscin channels were subsequently opened in Fiji and compressed into maximum intensity projections. Pre-constructed Iba1 ROI sets were then overlaid onto lipofuscin Z-projections and mean intensity of lipofuscin within each Iba1-positive ROI was recorded. Lipofuscin intensity data were analyzed with a linear effects model (LMEM) with mouse as a random effect. LMEMs were constructed in R using the lme4 and MASS libraries.

### Lipidomics

For lipidomic analysis, weighed tissue was homogenized in with ice-cold phosphate-buffered saline using a bead mill homogenizer. Tissue lysates (∼50 μg) were transferred to Pyrex glass tubes with a PTFE-liner cap. Lipids were then extracted by the Folch method (*54*). Briefly, 6 mL of ice-cold chloroform, methanol (2:1 v/v), and 1.5 mL of water were added to the samples and tubes were vortexed thoroughly to mix the samples homogenously with a polar and non-polar solvent. SPLASH LIPIDOMIX internal standards (Avanti Polar Lipids) were spiked in before the extraction. The organic phase of each sample was normalized by internal standards. After vortexing, samples were centrifuged for 20 min at 1000 rpm at 4°C to separate the organic and inorganic phases. Using a sterile glass pipette, the lower organic phase was transferred into a new glass tube, taking care to avoid the intermediate layer of cellular debris and precipitated proteins. The samples were dried under nitrogen flow until the solvents were completely dried. Samples were resuspended in 150 μL of chloroform: methanol 2:1 and stored R -80°C until mass spectrometer (MS) analysis.

#### Ganglioside extraction

∼100 μg homogenized tissues were extracted with methanol and spun down at 1000 x *g* for 20 mins to pellet proteins. Solvent layer containing gangliosides and other metabolites was collected in a fresh glass tube and dried under N2 stream. Dried extract was reconstituted in 1 mL LC-MS grade water and desalted by SOLA HRP SPE 30mg/2mL 96 well plate 1EA (Thermo Scientific #60509-001). Desalting cartridges were cleaned 3 times with 1mL methanol and equilibrated 3 times with LC-MS grade water. Next, the extracts dissolved in water were loaded onto the cartridge, washed with 2mL of water then gangliosides were eluted with 3 mL methanol. Eluate was dried under nitrogen flow and reconstituted in 300:150:50 of MeOH: water: chloroform.

#### LC-MS analysis of lipidomics

Lipids were separated using ultra-high-performance liquid chromatography (UHPLC) coupled with andem mass spectrometry. Briefly, UHPLC analysis was performed on a C30 reverse-phase column (Thermo Acclaim C30, 2.1 x 150 mm, 2.6 μm operated at 50°C; Thermo Fisher Scientific) connected to Vanquish Horizon UHPLC system and an Orbitrap Exploris 240 MS (Thermo Fisher Scientific) equipped with a heated electrospray ionization probe. 2 μL of each sample was analyzed separately, using positive and negative ionization modes. Mobile phase contained 40:60 water:acetonitrile (v:v), 10 mM ammonium formate, and 0.1% formic acid, and mobile phase B consisted of 90:10 2-propanol:acetonitrile, also including 10 mM ammonium formate, and 0.1% formic acid.

#### Chromatographic gradient

isocratic elution at 30% B from -3−0 minutes; linear increase from 30-43% B from 0-2 minutes; linear increase from 43-55% B from 2-2.1 minutes; linear increase from 55-65% B from 2.1-12 minutes; linear increase from 65-85% B from 12-18 minutes; linear increase from 85-100% B from 18-20 minutes; hold at 100-100% B from 20-25 minutes; linear decrease from 100-30% B from 25-25.1 minutes; hold at 30% B from 25.1-28 minutes. Flow rate, 0.26 mL/minute. Injection volume, 2 μL. Column temperature, 55°C.

#### Mass spectrometer parameters

ion transfer tube temperature, 300°C; vaporizer temperature, 275°C; Orbitrap resolution MS1, 120,000, MS2, 30,000; RF lens, 70%; maximum injection time 50 ms, MS2, 54 ms; AGC target MS1, standard, MS2, standard; positive ion voltage, 3250 V; negative ion voltage, 2500 V; Aux gas, 10 units; sheath gas, 40 units; sweep gas, 1 unit. HCD fragmentation, stepped 15%, 25%, 35%; data-dependent tandem mass spectrometry (ddMS2) cycle time, 1.5 s; isolation window, 1 m/z; microscans, 1 unit; intensity threshold, 1.0e4; dynamic exclusion time, 2.5 s; isotope exclusion, enable. Full scan mode with ddMS2 at m/z 250-1700 was performed. EASYICTM was used for internal calibration. The raw data were search and aligned using LipidSearch 5.0 with the precursor tolerance at 5 ppm and product tolerance at 8 ppm, and retention time deviation tolerance of 0.5 min. Further data post-processing filtering and normalization were performed using an in-house developed app, Lipidcruncher. All semi-targeted quantifications were done using area under the curve was normalized to the area under the curve for respective internal standards and amount of proteome/wet weight of tissue used.

#### LC-MS analysis of gangliosides

For gangliosides analysis, samples were analyzed on Vanquish UHPLC (Thermo, Germany) coupled to an Orbitrap Exploris 240 mass spectrometer (Thermo, Germany). Extracts were separated on Kinetex HILIC column (Phenomenex CAT 00D-4461-AN, 2.6 µm,100 x 2.1 mm). Solvent A was acetonitrile with 0.2% (v/v) acetic acid, and solvent B was water with 10 mM ammonium acetate, pH 6.1, adjusted with acetic acid. Column temperature was maintained at 50°C. LC gradient started with constant flow rate of 0.6 mL/min with 12.3% B at 0 min, linear gradient of 12.3% B to 22.1% B from 1 min to 15 mins. Column was equilibrated between runs by 12.3% B for 5 mins. The spray voltage was set to −2.5 kV, and the heated capillary and the HESI were held at 300°C and 250°C, respectively. The S-lens RF level was set to 50, and the sheath and auxiliary gas were set to 40 and 5 units, respectively. These conditions were held constant during the acquisitions. All measurements were performed in semi-targeted semi-quantitative mode. The raw data was performed search and aligned with Lipidsearch 5.0. Further data post-processing filtering and normalization were performed using an in-house developed app, Lipidcruncher.

### Plasma neurofilament light chain

Plasma neurofilament light chain (NfL) levels were analyzed using the Quanterix Simoa NF-light V2 Advantage Kit. Prior to dilution, samples were spun 5 minutes at 10,000 x *g* at 4°C to remove any debris. Samples were diluted 1:4 in sample diluent and were plated in duplicate along with kit calibrators. Simoa detector reagent and bead reagent were added to samples and were incubated for 30 minutes at 30°C on a shaker at 800 rpm. Next, samples were washed with Simoa Wash Buffer A using the Simoa Microplate Washer and the Quanterix 2-step protocol. SBG reagent was added and samples were allowed to incubate 10 minutes at 30°C on the shaker at 800 rpm. The sample plate was then washed using the 2-step washing protocol where the beads bound to sample were resuspended twice in Simao Wash Buffer B before aspiration of buffer. The plate was allowed to dry for 10 minutes at room temperature prior to loading in the Quanterix SR-X instrument where NfL concentration was determined by interpolation against the calibration curve.

### Statistics

All statistical analysis was performed in GraphPad Prism Versions 9 and 10 with the following exceptions. For Microglial lipofuscinosis data LMEM analyses were performed in R using the lme4 and MASS libraries. The lipidomic dataset was analyzed using Metaboanalyst 5.0 one factor statically analysis. Data in bar graphs are presented as ± SEM with individual data points.

DAB immunohistochemistry experiments for SCMAS, CD68, and Galectin-3 were imaged across multiple regions and were analyzed using mixed-effects analyses. Autofluorescence was also imaged across multiple regions and was analyzed using mixed-effect analysis. For data comparing all three viral vectors in terms of individual outcome measures we used the Brown-Forsythe test of variances to test the assumption of homogeneity of variance across treatment groups. In cases where this assumption was violated, primarily due to many zero values in the GFP negative control group, we employed the Brown-Forsythe variant of the one-way ANOVA with pairwise comparisons using Tamhane test to account for the resulting heteroscedasticity. In all other cases, we used an ordinary one-way ANOVA. Microglial lipofuscinosis data were analyzed with a linear effect model (LMEM) with mouse as a random effect. Lipidomics data were analyzed using Metaboanalyst 5.0 one factor statistical analysis. When using all lipids in the dataset for volcano plots, data were first filtered using interquartile range (IQR) of 25%. For analysis of just BMP and ganglioside species shown in heat maps, no filtering was necessary. To compare lipidomic signatures of AAV-GFP treated *Grn*^−/−^ and *Grn^+^*^/+^ mice, volcano plots were generated from –Log10(*p*-value) and Fold change of 1.5. For heatmaps with all 4 groups, ANOVAs were run with parameters set to FDR *p*-value of 0.05 with Tukey’s post *hoc* analysis. For graphs with percent correction for BMP/Ganglioside species, only lipids that were significantly different and ±1.5 log2-fold change in *Grn*^−/−^ vs *Grn*^+/+^ mice were used. Percent correction was calculated by subtracting knockout levels from experimental (NT or CT) and dividing that number by wild-type minus knockout, setting the graph axis to 0% for knockout mice and 100% for wild-type mice (experimental group difference from *Grn*^−/−^:GFP divided by *Grn*^+/+^:GFP difference from *Grn*^−/−^ :GFP). Comparisons of the most abundant lipid species were done with one-Way ANOVA. Plasma NfL data were analyzed using a one-way ANOVA.

## List of Supplementary Materials

Supplementary Fig 1.

Supplementary Fig. 2.

Supplementary Fig. 3.

## Funding

This work was supported by the

Bluefield Project to Cure FTD (EDR, TCW, RVF)

National Institutes of Health grant F30AG071114 (SNK)

Howard Hughes Medical Institute (TCW)

## Author contributions

Conceptualization: EDR

Funding acquisition: EDR, SNK, TCW, RVF

Methodology: SNK, SNF, YAA, TCW, RVF, AEA, EDR

Investigation: SNK, SNF, KIW, YAA, AEA

Formal analysis: SNK, SNF, YAA, CFM

Writing – original draft: SNK, EDR

Writing – review & editing: SNK, SNF, CFM, YAA, TCW, RVF, AEA, EDR

## Data and materials availability

All data needed to evaluate the conclusions in the paper are present in the paper and/or the Supplementary Materials.

