## Supplemental Material for "Carboxy-terminal blockade of sortilin binding enhances progranulin gene therapy, a potential treatment for frontotemporal dementia"

### SUPPLEMENTAL FIGURES

Fig S1

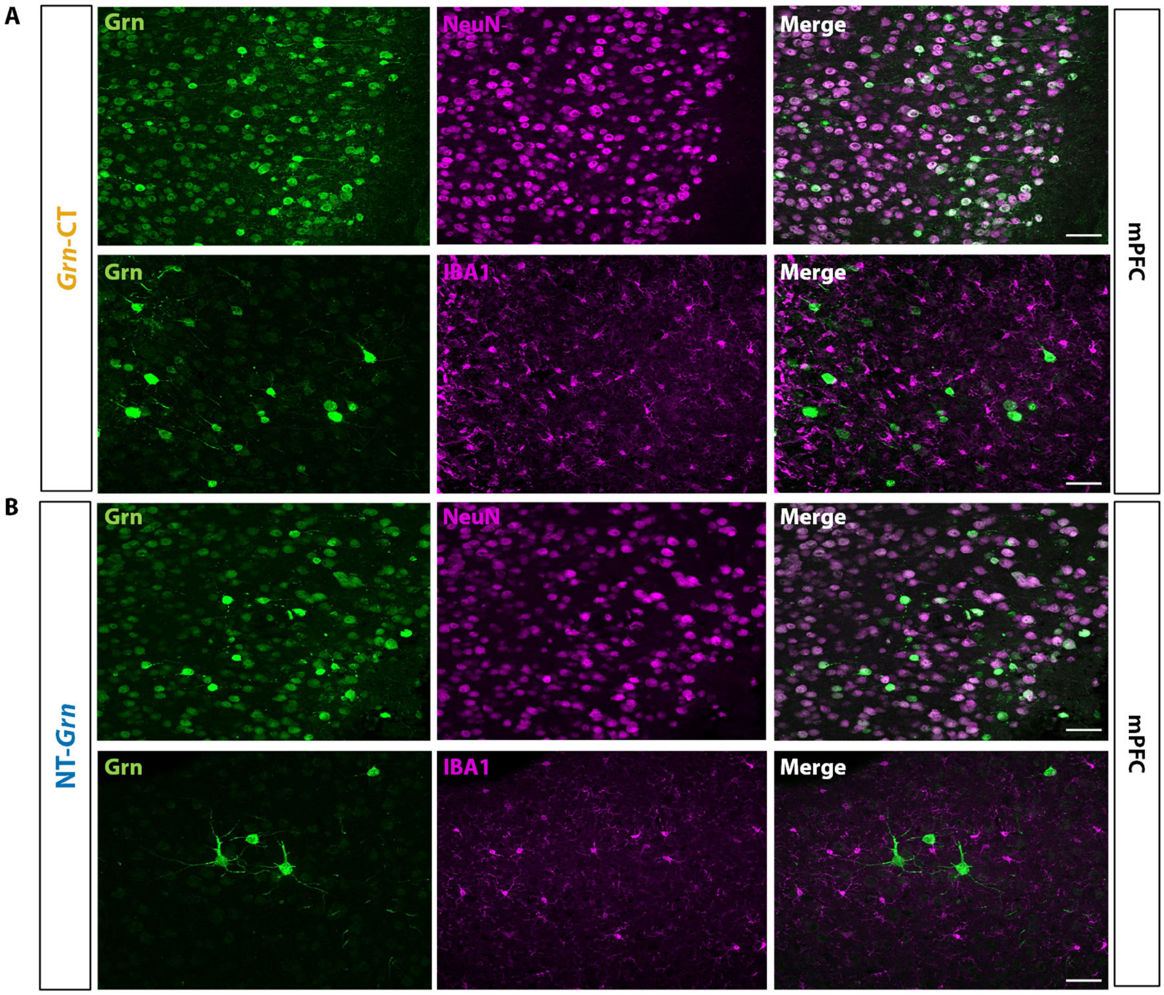


**Fig S1. Progranulin derived from both *Grn*-CT and NT-*Grn* is found primarily in neurons, not microglia.** (**A**) Representative images of mPFC from mice treated with *Grn*-CT. Progranulin primarily co-labeled with NeuN (top), not Iba1 (bottom). (**B**) Representative images of mPFC from mice treated with NT-*Grn*. Similarly, progranulin primarily co-labeled with NeuN (top), not Iba1 (bottom).

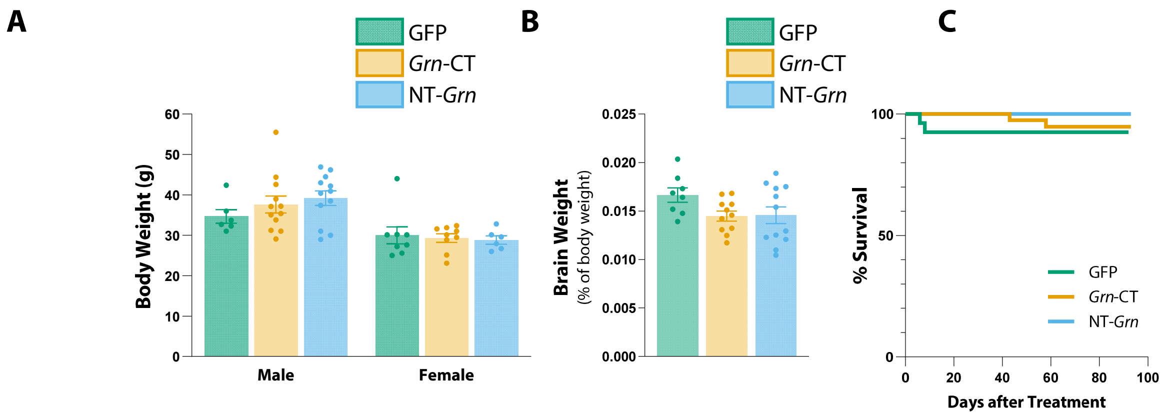


**Fig S2.** No differences in body weight, brain weight or survival with *Grn*-CT or NT-*Grn* treatment relative to GFP treatment**.** (**A**) Treatment with *Grn*-CT or NT-*Grn* does not precipitate measurable changes in body weight relative to GFP. Restricted mixed-effects model (REML) ($P < 0.0001$ for sex effect, $P = 0.7090$ for AAV effect), Tukey's MC test; GFP vs. *Grn*-CT ($P = 0.8315$), GFP vs. NT-*Grn* ($P = 0.6921$), *Grn*-CT vs. NT-*Grn* ($P = 0.9517$)). (**B**) Treatment with *Grn*-CT or NT-*Grn* does not precipitate measurable changes in brain weight relative to GFP. One way ANOVA ($P = 0.1104$); Tukey's MC test; GFP vs. *Grn*-CT ($P = 0.1389$), GFP vs. NT-*Grn* ($P = 0.1500$), *Grn*-CT vs. NT-*Grn* ($P = 0.9961$); *n*=8–12 mice. (**C**) Treatment with *Grn*-CT or NT-*Grn* does not change survival relative to GFP. Log-rank /Mantel Cox test; $P = 0.2987$; *n*=27 mice in GFP group, 40 mice in *Grn*-CT group, 35 mice in NT-*Grn* group.

Fig S3

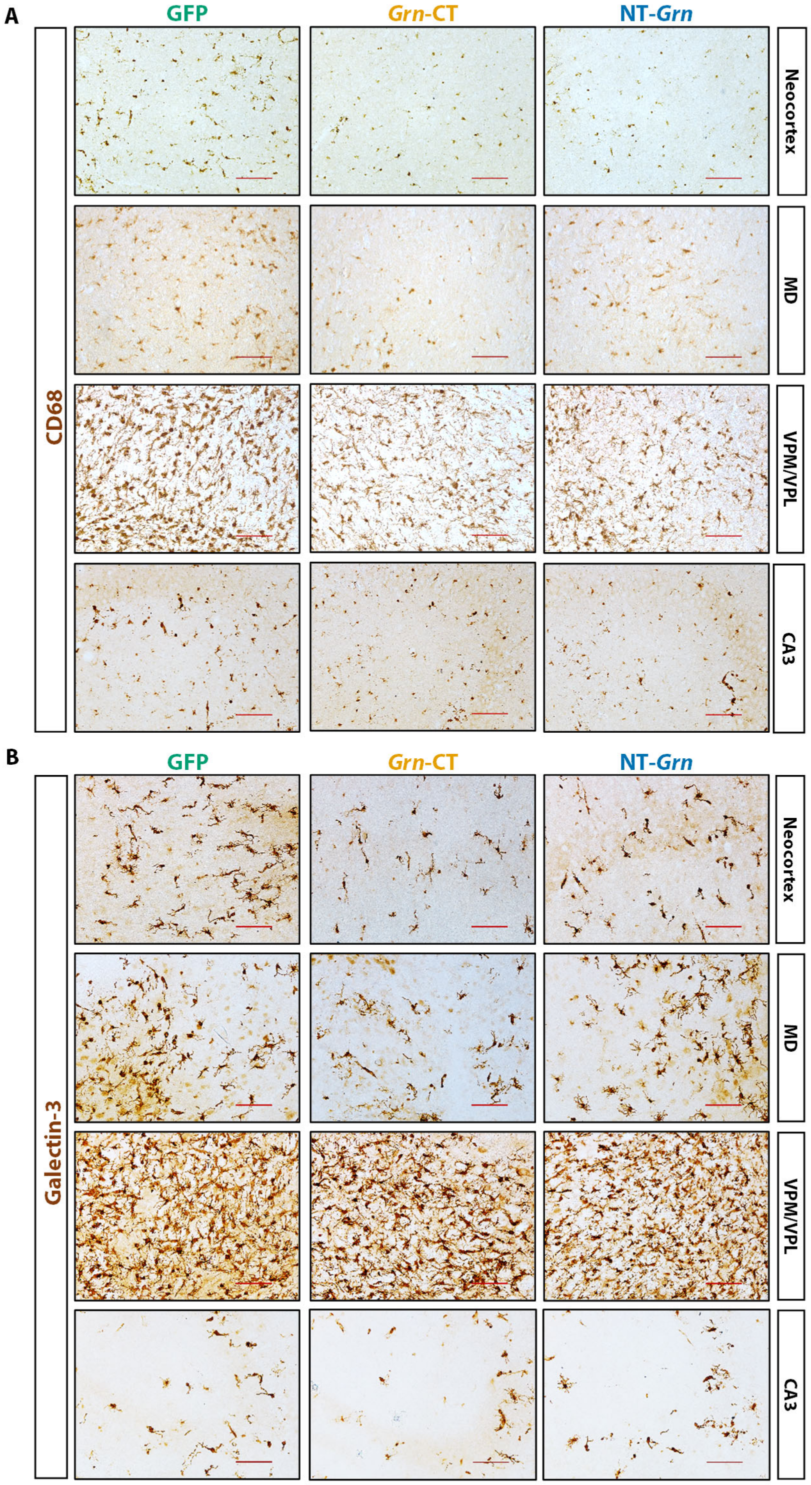


**Fig S3.** Representative micrographs of **(A)** CD68 and **(B)** Galectin staining in other brains regions quantified in Figure 3. Scale bars, 100µm.
